# GO Big or Go Home: A New Gene Ontology Subset that Improves Plant Gene Function Prediction

**DOI:** 10.1101/2025.01.26.634938

**Authors:** Leila Fattel, Carolyn J. Lawrence-Dill

**Affiliations:** Interdepartmental Genetics and Genomics, Iowa State University, Ames, Iowa, 50011, USA; Department of Agronomy, Iowa State University, Ames, Iowa, 50011, USA; Department of Soil and Crop Sciences, Colorado State University, Fort Collins, Colorado, 80523, USA

**Author notes:** Contributing authors.

**Keywords:** Gene Ontology, Plants, Gene Function, Annotation

## Abstract

**Background:** The availability of gene function prediction datasets helps researchers to consider possible functions for uncharacterized genes for hypothesis generation, candidate gene prioritization, and many other applications. Many such datasets are based on the Gene Ontology (GO) function graph. For plants this can be problematic because the most specific GO terms available are often derived from the biology of non-plant taxa (e.g., functions specific to nerve function would not seem likely to map to plant biological processes given that plants lack nerves). To balance the need for functional specificity while limiting to functions relevant to plant biology, researchers often limit to the GO Slim plant subset, but, by design, that subset consists of very general terms and limits real utility for e.g., specific hypothesis generation. Worse yet, sometimes researchers choose to simply throw out terms if they are not relevant to plant biology (rather than traversing the GO graph to select the most specific term in that hierarchy that is compatible with plant biology).

**Results:** We created GO Big, a Gene Ontology subset type, to improve the biological relevance of gene function predictions for taxon-specific biology applications. GO Big plant subsets retain maximal functional specificity for hypothesis generation while limiting to terms applicable to the biology of plants. In brief, we used a curatorial approach to generate two GO Big subsets, a general subset derived from terms with experimentally validated functions across Viridiplantae species, and a species-specific subset for maize (*Zea mays* ssp. *mays*).

**Conclusion:** Annotating genes with assignments that better reflect the biology of a taxon can pave the way for more biologically accurate and testable hypotheses for genes of interest. The subsets produced here can help plant biologists limit genome-wide gene function prediction sets to functions possible for plant genes, and the process to generate GO Big subsets is described in detail to enable others to create GO Big subsets for additional taxon sets, including ones for protists, fungi, and other phylogenetic categories.

## 1 Introduction

The development of the bio-ontologies has been integral for the structured representation of biological knowledge to enable computational approaches [1]. Of note is the regularly updated Gene Ontology (GO) created and maintained by the Gene Ontology Consortium [2, 3]. The GO is a directed acyclic graph (DAG) of standardized vocabulary terms that represents the functional characteristics of genes and gene products [4]. GO has been used by researchers to predict the function of a gene of interest, to find common functions between genes located in one or more organisms, to evaluate the abundant functions in gene expression studies, and to hypothesize possible protein-protein interactions, to name a few [5]. There are three categories of GO terms: Biological Process (BP), Molecular Function (MF), and Cellular Component (CC) [2]. The terms (nodes) in the GO knowledgebase are connected through relations (edges), such as “is a”, “part of”, “regulates”, “positively regulates”, “negatively regulates”, and more [3]. In addition, GO annotation assignments are supported by evidence codes so that the logic for how a term was assigned is explained. These evidence codes are classified under six general categories: experimental, phylogenetic, computational, author statements, curatorial statements, and automatically generated annotations [3]. Initially, GO terms were derived from three model organism databases [2]: Saccharomyces Genome Database (SGD) [6], FlyBase [7], and Mouse Genome Informatics (MGI) [8]. Since then, several collaborators from various model organism and protein databases have joined in the efforts of the GO consortium to develop and expand the framework to include several animal, plant, fungal, bacterial, and viral gene function annotations [3, 9]. As of April 2024, there are 42,255 GO terms in use [10].

Gene function prediction tools provide researchers with gene function assignments that can help them to identify genes responsible for phenotypes of interest, to discover potential drug targets, and to find underlying causes of diseases [11]. An example of a plant-specific gene function prediction pipeline is the Gene Ontology Meta Annotator for Plants (GOMAP), which produces high-coverage and reproducible GO-based functional annotation datasets [12]. However, a problem for gene function prediction in plants is the lack of some specific GO terms related to plant function, coupled with the presence of some very detailed gene function term assignments derived from animal, fungi, and microorganisms. This results in less accurate annotation assignments for plants [13]. For example, a study we conducted discussed that some plant genes were assigned with GO terms like “animal organ development” (GO:0048513), while also being annotated with plant-specific functions [14].

GO slims are reduced and manageable subsets of the GO that contain terms describing gene functions in a broad manner [9, 15]. Several GO slims currently exist and can be found in the GO database (https://geneontology.org/docs/downloadontology/#subsets) [16]. Subsets can be species-specific, such as that for the fungus *Candida albicans* (GO Slim candida) or even kingdom-specific, such as that for plants (GO Slim plant) [5]. These high-level GO terms are tagged within the GO DAG to allow the display of (and functional limitation to) each subset to which the term belongs [3]. The main purpose of a GO slim is to represent and map gene function annotations to a summary of general biological functions [17]. The small number of high-level GO terms that are members of each GO Slim subset were selected as a set of terms that would be most effective at capturing all categories of functions derived from all annotated genes of the taxa. GO slims are often used in gene expression studies to highlight over- and under-represented functions shared by a group of genes in one or more organism [18].

When it comes to gene function prediction, annotating with a GO term that is as specific as possible, while allowing for incorrect assignments, can be beneficial as these annotations can become starting points for generating and testing hypotheses for genes of unknown functions. However, some plant biology researchers are only interested in plant-possible functions. So far, there had been two GO-based approaches to help with that: 1) mapping against the plant GO slim subset to get higher-level, biologically relevant functions, and 2) relying on the taxon constraints set by the GO Consortium through “only in taxon” and “never in taxon” relationships allocated to some GO terms, a quality control method used by curators to detect incorrect gene function annotations [19]. Here, we discuss a third approach that is evidence-based and dependent on the taxa of interest. The GO Big subsets maximize taxonomic unit specificity while limiting to terms relevant to the biology of the included taxa to assist with automated gene function prediction analyses. We also describe the methods used to create a species-specific subset (e.g., GO Big maize) and a more general green plants subset (GO Big plant).

## 2 Methods

Creating GO subset files can be time- and labor-intensive. In this section, we demonstrate creating the GO Big subsets, “GO Big maize” and “GO Big plant”, using two methods. They both consist of the following steps: accumulating GO terms assigned with experimental evidence codes, collecting the ancestors of these GO terms, tagging the GO terms of interest as members of the GO Big subset, and creating the GO Big subset file.

### 2.1 The GO Big maize subset (as a species-specific example)

GO terms assigned through manual curation from experimental literature of the *Zea mays* ssp. *mays* genome were gathered using the Gene Ontology Annotation (GOA) database [20]. The filters were chosen as follows:

- Taxon: *Zea mays* (taxon ID: 4577) and exact match.
- GO terms: For each GO category at a time, BP, MF, and CC (selected under the “Aspect” filter), their respective GO terms belonging to the GO Slim plant subset were added as identifiers (BP: 45; MF: 25; CC:24), and the option “use these terms as a GO slim” was selected to find GO terms that map back to the plant subset. Alternatively, one can select all GO categories under the “Aspect” filter at the same time and take advantage of the “select terms from GO slim” option under “GO terms” by choosing “goslim plant”. However, this was not done here because the output files can be too large and we settled on splitting them by GO aspect instead.
- Evidence: To avoid collecting plant genes with errant function assignments, only experimental evidence codes were selected. Experimental evidence codes indicate that there is direct evidence within the literature that supports the annotation of a gene. The experimental evidence codes - along with their child terms - chosen were ECO:0000352 (evidence used in manual assertion), ECO:0000269 (experimental evidence used in manual assertion), ECO:0000314 (direct assay evidence used in manual assertion), ECO:0000315 (mutant phenotype evidence used in manual assertion), ECO:0000316 (genetic interaction evidence used in manual assertion), ECO:0000353 (physical interaction evidence used in manual assertion), ECO:0000270 (expression pattern evidence used in manual assertion), ECO:0006056 (high throughput evidence used in manual assertion), ECO:0007005 (high throughput direct assay evidence used in manual assertion), ECO:0007001 (high throughput mutant phenotypic evidence used in manual assertion), ECO:0007003 (high throughput genetic interaction phenotypic evidence used in manual assertion), and ECO:0007007 (high throughout expression pattern evidence used in manual assertion).

Once the filters were applied, the files were downloaded and the annotations under the “GO term” column were looked up using QuickGO [21] for manual collection of these terms and their ancestors all the way up to the root term. While collecting all these terms, their descriptions were inspected to make sure they had functions applicable in the context of plants. After GO term collection, Protégé [22] was used to tag these terms as members of the GO Big maize subset using the go.owl file release “2023-10-09”. Robot [23] was then used to create the gobig maize.owl file (Web Ontology Language - a file format for knowledge representations like ontologies) by selecting and filtering for the terms tagged as part of the subset. Subsequently, the gobig maize.obo file (Open Biomedical Ontologies - a file format for representing ontologies and controlled vocabularies) was also created.

### 2.2 The GO Big plant subset

To create the GO Big subset for green plants (Viridiplantae), we included more species to capture diverse plant functions. All GO annotations with experimental evidence codes assigned to plant species that are descendants of Viridiplantae (taxon ID: 33090) were collected from the GOA database. Unlike the GO Big maize subset, we did not filter the GO terms using the GO Slim plant subset as identifiers so that all GO annotations with manually assigned evidence codes would be captured.

The generation of the GO Big plant subset (gobig plant.owl and gobig plant.obo files) was similar to the method described for GO Big maize. To automate the collection of the GO ancestors, we created python scripts using the GOATOOLS package [24]. Here, the go.owl file release “2024-04-24” was used to manually tag the terms belonging to the GO Big plant subset in Protégé. Additionally, using the go taxon constraints.owl file provided by the GO Consortium [25], any GO term labelled with “only in taxon” for the class Viridiplantae that was not experimentally assigned to a plant gene was added to the subset (as well as its ancestor terms) as a way to further capture more plant functions in this subset. The method is summarized in Figure 1. All the files and scripts described here can be found in our GitHub repository [26].

**Fig. 1.**
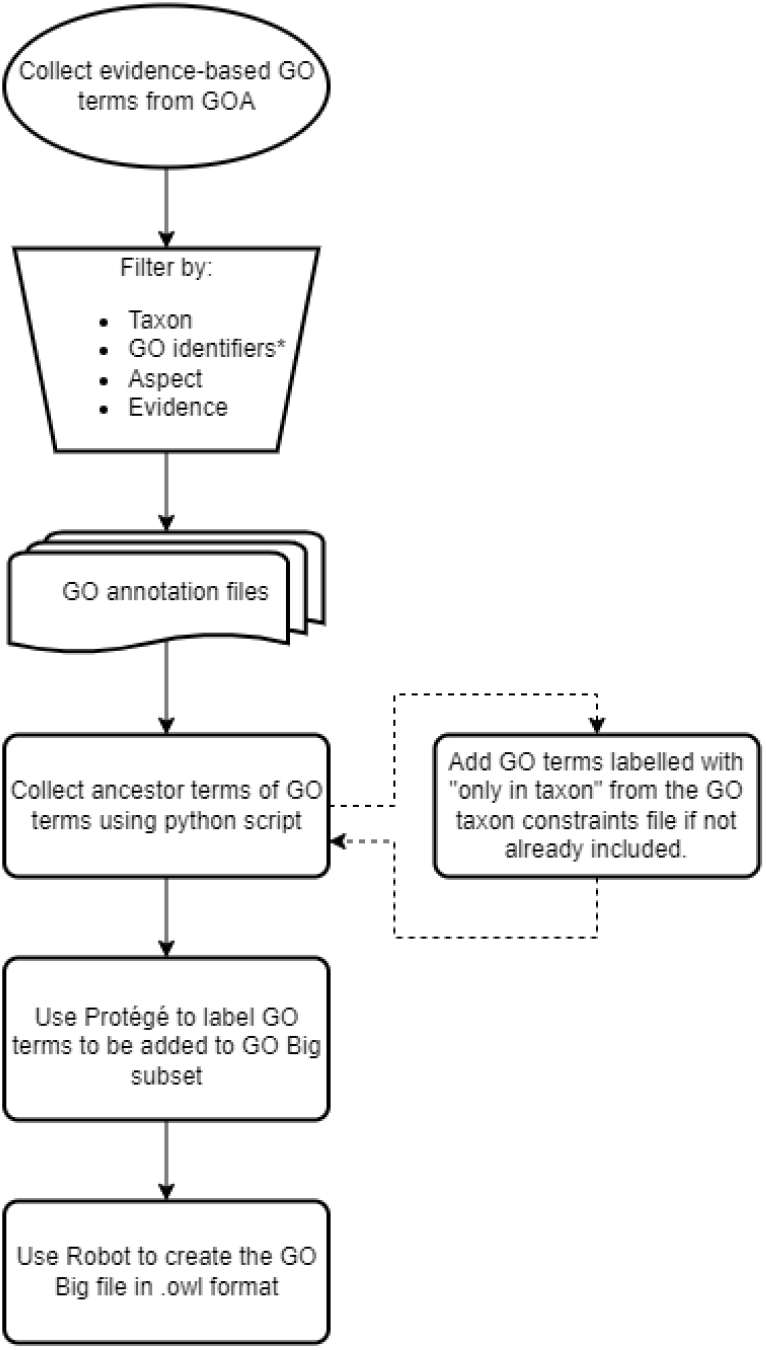
Workflow for creating a GO Big subset. Oval represents start, inverted trapezoid represents a manual process, curved rectangle represents multiple documents, and rectangles represent tasks with outputs. * symbolizes only in GO Big maize. Dashed arrows represent the additional process included for GO Big plants.

## 3 Results

### 3.1 The GO Big subsets

Because the GO Big plant subset includes GO terms derived from more than one plant, it is (approximately 11X) larger than the GO Big maize subset, as seen in Table 1. The GO Big plant subset has a total of 10,611 GO terms that includes 6,236 BP, 1,087 CC, and 3,288 MF terms. Meanwhile, the GO Big maize subset has a total of 1,003 GO terms that contain 599 BP, 88 CC, and 316 MF terms.

**Table 1.** Number of GO terms by GO aspect in each GO set.

| GO Set | GO Aspect [count (*)] |  |  |  |
| --- | --- | --- | --- | --- |
|  | Biological Process | Cellular Component | Molecular Function | All |
| GO Big plant | 6,236 (58.77%) | 1,087 (10.24%) | 3,288 (30.99%) | 10,611 |
| GO Big maize | 599 (59.72%) | 88 (8.77%) | 316 (31.51%) | 1,003 |
| GO Slim plant | 46 (47.42%) | 25 (25.78%) | 26 (26.80%) | 97 |
| GO release "2023-10" | 27,538 (64.28%) | 4,060 (9.48%) | 11,239 (26.24%) | 42,837 |
| GO release "2024-04" | 27,047 (64.01%) | 4,057 (9.60%) | 11,151 (26.39%) | 42,255 |

\*Parenthetical numbers reflect percentage of terms by aspect out of all terms available in the respective set.

### 3.2 General comparison of the GO Big subsets with the full GO DAG and GO Slim plant subset

Both the GO Big subsets contain more terms than the GO Slim plant subset. Although the GO Big maize subset (1,003 GO terms) includes terms derived from experiment-based annotations from only one plant, it is still approximately 10X larger than the GO Slim plant subset (97 GO terms). At the same time, the GO Big plant subset (10,611) contains around 109X the GO terms included in GO Slim plant. On the other hand, GO Big maize and GO Big plant only cover ∼2% and ∼25% of the entire GO corpus, respectively. A full comparison of the number of GO terms by aspect is shown in Table 1. To demonstrate the impact of interpreting gene function based on GO Big on functional annotation, we provide an example in the next section.

### 3.3 Demonstration: non-plant annotations for a plant gene

The SET (Su(var)3-9, Enhancer-of-zeste and Trithorax) domain is often found in various histone methyltransferases that play a role in epigenetic modifications [27]. In plants, histone methyltransferases play a role in several biological processes, including morphogenesis, hormone regulation, flowering time, shoot and root branching, seed development, circadian cycle, and some stress responses [27, 28]. A recent study reported that the SET domain gene 130 (*sdg130* ) is a candidate gene involved in the regulation of vegetative phase change in maize upon jasmonic acid treatment [29]. The GO annotations for *sdg130* (gene ID Zm00001eb216690) in the maize B73v5 genome [30] using the Gene Ontology Meta Annotator for Plants (GOMAP) [12] are shown in Table 2. The annotations mainly describe functions relating to histone methylation and transcription processes. However, of the 28 GO terms, four describe non-plant functions: “skeletal muscle myosin thick filament assembly” (GO:0030241), “sarcomerogenesis” (GO:0048769), “cardiac myofibril assembly” (GO:0055003), and “skeletal muscle organ development” (GO:0060538). A common approach to deal with terms describing non-plant functions is to ignore them. However, these errant GO annotations may be useful.

**Table 2.**
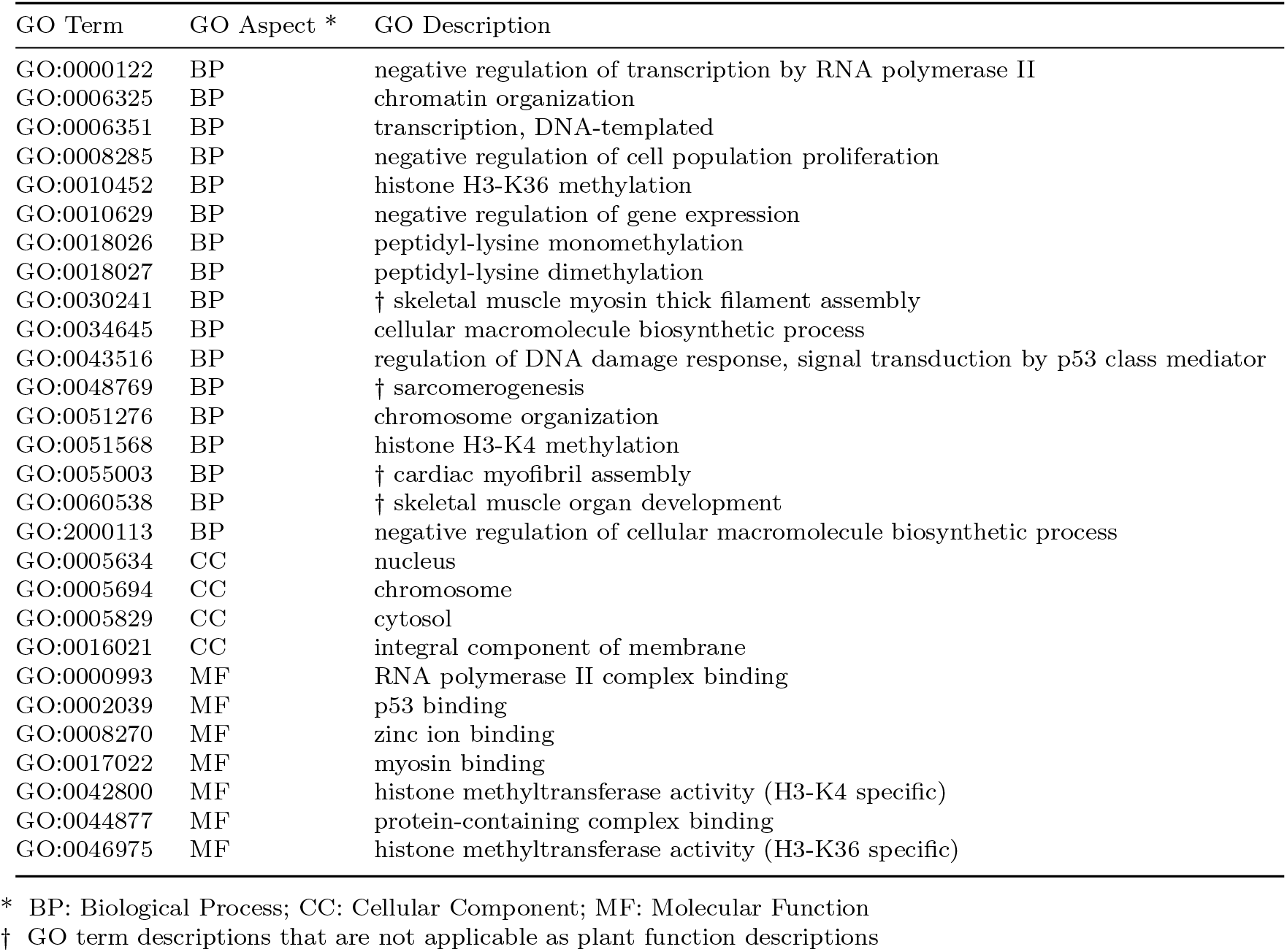
GO term annotations for the maize B73v5 *sdg130* gene assigned by GOMAP.

| GO Term | GO Aspect * | GO Description |
| --- | --- | --- |
| GO:0000122 | BP | negative regulation of transcription by RNA polymerase II |
| GO:0006325 | BP | chromatin organization |
| GO:0006351 | BP | transcription, DNA-templated |
| GO:0008285 | BP | negative regulation of cell population proliferation |
| GO:0010452 | BP | histone H3-K36 methylation |
| GO:0010629 | BP | negative regulation of gene expression |
| GO:0018026 | BP | peptidyl-lysine monomethylation |
| GO:0018027 | BP | peptidyl-lysine dimethylation |
| GO:0030241 | BP | † skeletal muscle myosin thick filament assembly |
| GO:0034645 | BP | cellular macromolecule biosynthetic process |
| GO:0043516 | BP | regulation of DNA damage response, signal transduction by p53 class mediator |
| GO:0048769 | BP | † sarcomerogenesis |
| GO:0051276 | BP | chromosome organization |
| GO:0051568 | BP | histone H3-K4 methylation |
| GO:0055003 | BP | † cardiac myofibril assembly |
| GO:0060538 | BP | † skeletal muscle organ development |
| GO:2000113 | BP | negative regulation of cellular macromolecule biosynthetic process |
| GO:0005634 | CC | nucleus |
| GO:0005694 | CC | chromosome |
| GO:0005829 | CC | cytosol |
| GO:0016021 | CC | integral component of membrane |
| GO:0000993 | MF | RNA polymerase II complex binding |
| GO:0002039 | MF | p53 binding |
| GO:0008270 | MF | zinc ion binding |
| GO:0017022 | MF | myosin binding |
| GO:0042800 | MF | histone methyltransferase activity (H3-K4 specific) |
| GO:0044877 | MF | protein-containing complex binding |
| GO:0046975 | MF | histone methyltransferase activity (H3-K36 specific) |

\* BP: Biological Process; CC: Cellular Component; MF: Molecular Function
† GO term descriptions that are not applicable as plant function descriptions

To evaluate if errant GO terms can provide useful insights while staying relevant to plant biology, we mapped the 28 GO annotations to the GO Slim plant, GO Big maize, and GO Big plant subsets using GOTermMapper [9, 31] to identify information loss in each subset (see Additional File 1). Mapping resulted in 21 GO terms (13 BP, 4 MF, and 4 CC) using the GO Slim plant subset, 103 GO terms (75 BP, 16 MF, and 12 CC) using the GO Big maize subset, and 149 GO terms (105 BP, 32 MF, and 12 CC) using the GO Big plant subset. In general, the terms from the GO Slim plant subset describe high-level processes like “protein metabolic process” (GO:0019538) and “multicellular organism development” (GO:0007275), and functions like “protein binding” (GO:0005515) and “transferase activity” (GO:0016740). With the GO Big maize subset, however, the GO term descriptions start becoming more specific, and include processes relating to transcription (“regulation of DNA-templated transcription” (GO:0006355); “chromosome organization” (GO:0051276); “gene expression” (GO:0010467)), and morphogenesis (“anatomical structure morphogenesis” (GO:0009653); “system development” (GO:0048731); “tissue development” (GO:0009888)). Both of which are biological processes that *sdg130* has been reported to be involved in as described previously. In addition, cytoskeleton-related GO terms, including “cytoskeleton organization” (GO:0007010) and “actin cytoskeleton 8 organization” (GO:0030036) were detected, both of which are ancestor terms for “sarcomerogenesis” (GO:0048769), one of the errant GO annotations. As expected, using the GO Big plant subset resulted in even more specific functions, among which that describe histone methylation, such as “histone H3 methyltransferase activity” (GO:0140938) and “protein methylation” (GO:0006479), presenting more biologically meaningful predictions of the various roles of *sdg130* in *Z. mays* in comparison to the GO Slim plant subset, while also making good use of non-plant function descriptions without fully incorporating them.

For demonstration purposes, we mapped one of the errant GO terms, “sarcomerogenesis” (GO:0048769), to the GO Slim plant (Figure 2), GO Big maize (Figure 3), and GO Big plant (Figure 4) subsets to show the ancestor terms derived, as well as the depth (the length of the longest path from the top) of the most specific GO term in each case. In Figure 2, the GO Slim plant resulted in only four high-level GO terms and a maximum depth of three (GO:0016043 and GO:0030154) in the DAG. Although the ancestor terms obtained from the GO Slim plant subset do give a general idea that the possible functions of *sdg130* include structure development and cellular component organization, the GO Big maize subset (Figure 3) captured 13 ancestor GO terms that cover more processes relating to morphogenesis and, more specifically, the actin filament component of the cytoskeleton with a maximum depth of six (GO:0030036). Finally, the GO Big plant subset (Figure 4) had the highest coverage of a total of 22 ancestor GO terms, with a maximum depth of seven (GO:0031032).

**Fig. 2.**
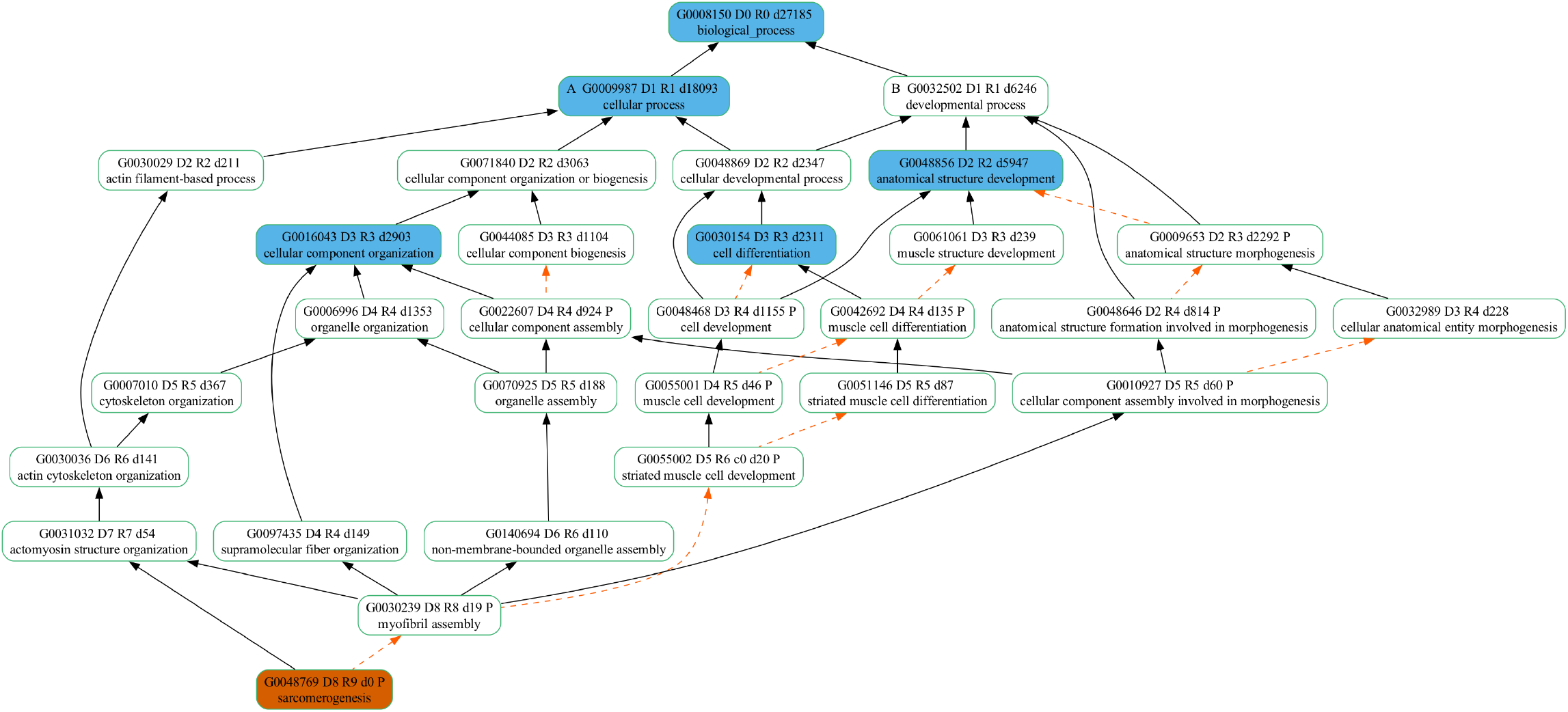
Mapping “sarcomerogenesis” to the GO Slim plant subset. Terms appear in oval boxes. Orange fill represents the GO term assigned. Blue fill represents GO terms included in the GO Slim plant subset. Relationships between terms are shown as arrows. Solid black arrows denote “is a” relationships, while dashed arrows in orange denote “part of” relationships. In each oval box, the first line is the GO term header. Term identifiers are formatted as G#######. Letters A and B prepending the term identifier demarcate the highest-level ancestors of the term. The letter D prepends a number indicating the maximum distance from the root term (biological process) to the child term using only “is a” relationships; R is the maximum distance from the root term to the child term using both “is a” and “part of” relationships; d is the descendants count.

**Fig. 3.**
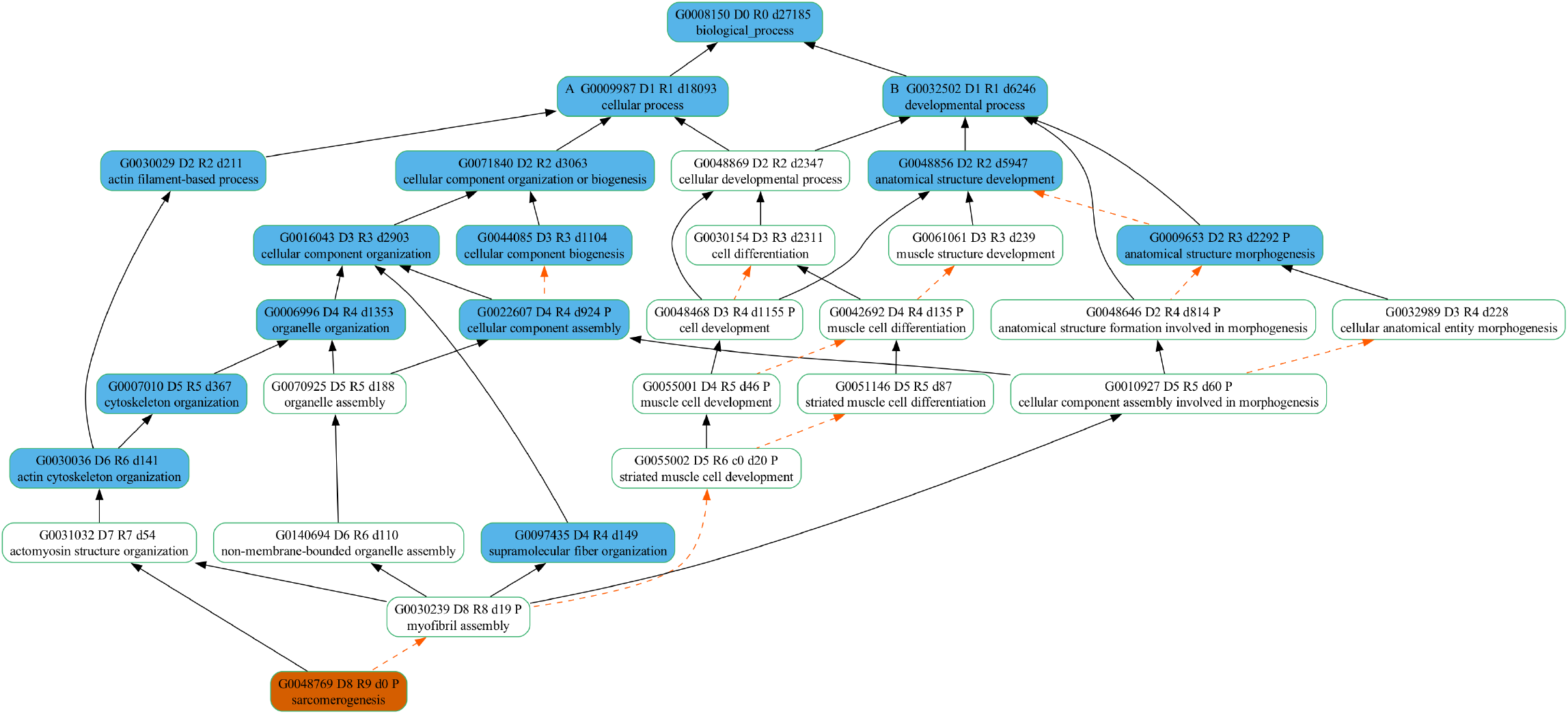
Mapping “sarcomerogenesis” to the GO Big maize subset. Terms appear in oval boxes. Orange fill represents the GO term assigned. Blue fill represents GO terms included in the GO Big maize subset. Solid black arrows denote “is a” relationships, while dashed arrows in orange denote “part of” relationships. In each oval box, the first line is the GO term header. Term identifiers are formatted as G#######. Letters A and B prepending the term identifier demarcate the highest-level ancestors of the term. The letter D prepends a number indicating the maximum distance from the root term (biological process) to the child term using only “is a” relationships; R is the maximum distance from the root term to the child term using both “is a” and “part of” relationships; d is the descendants count.

**Fig. 4.**
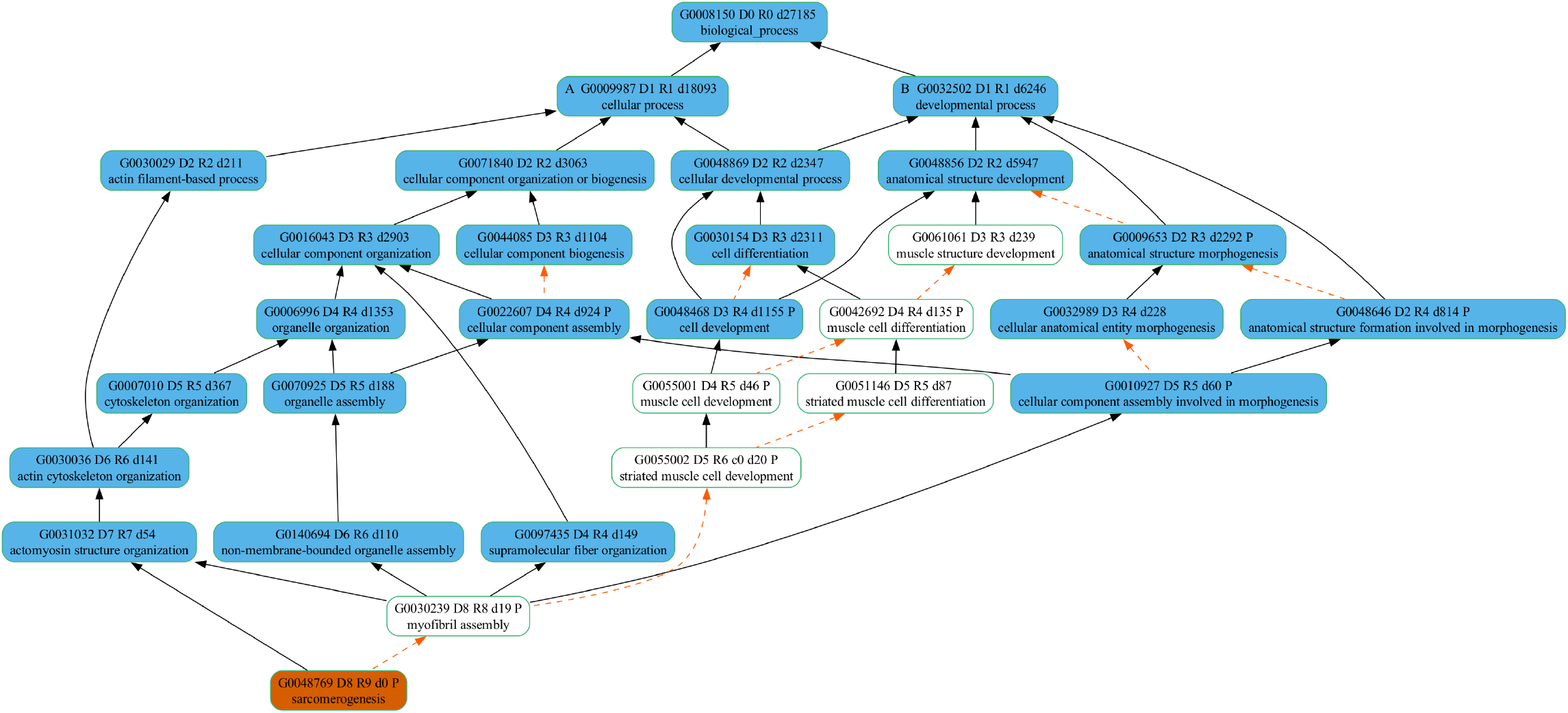
Mapping “sarcomerogenesis” to the GO Big plant subset. Terms appear in oval boxes. Orange fill represents the GO term assigned. Blue fill represents GO terms included in the GO Big plant subset. Solid black arrows denote “is a” relationships, while dashed arrows in orange denote “part of” relationships. In each oval box, the first line is the GO term header. Term identifiers are formatted as G#######. Letters A and B prepending the term identifier demarcate the highest-level ancestors of the term. The letter D prepends a number indicating the maximum distance from the root term (biological process) to the child term using only “is a” relationships; R is the maximum distance from the root term to the child term using both “is a” and “part of” relationships; d is the descendants count.

Although the GO Big plant subset covers more terms describing plant functions and processes than the GO Big maize subset, some errant terms were still somehow incorporated. Upon further analysis of the GO terms that resulted from mapping the 28 GO term annotations (Additional File 1), two terms from the GO Big plant subset stand out: “animal organ development” (GO:0048513) and “circulatory system development” (GO:0072359). To investigate the reason behind the incorporation of these terms (even when only selecting experiment-based evidence codes), we discuss the two examples below.

#### 3.3.1 Example 1: incorporation of GO terms of non-plant functions in a plant subset

To identify the cause of the incorporation of the non-plant function GO term “animal organ development” (GO:0048513), which indeed has not been experimentally annotated to any Viridiplantae gene in the GOA database, we inspected the presence of any of its child terms in the GO Big plant subset. Since the subset was built in a manner that preserves the structure of the GO DAG, any GO term added to the subset will have its ancestor terms included there. Indeed, the term “mesenchyme migration” (GO:0090131) has led to the integration of “animal organ development” (GO:0048513) in the GO Big plant subset as an ancestor term. The child term itself appears problematic in this context as the mesenchyme refers to a population of multipotent cells found in animal models [32]. Interestingly, this term has been manually assigned to the *Z. mays* gene *B4G1E1* (UniProtKB:B4G1E1). Further investigation in the literature has shown that there are studies that report plant products and bioactive compounds playing a role in mesenchymal stem cell proliferation and differentiation 9 [33, 34]. Although the GO term itself is not representative of a function that is present in plant physiology, it seems to reflect a function that represents regulatory interactions between plants and other species. This highlights the importance of inspecting seemingly errant GO terms assigned to plant gene products from a perspective that considers plant-non-plant interactions.

Another child term for the GO term “animal organ development” (GO:0048513) found in our GO Big plant subset is “interkinetic nuclear migration” (GO:0022027). The term “interkinetic nuclear migration” itself has been first used to describe nuclear movements observed during neural proliferation in vertebrates [35]. Although the term does not seem applicable in the context of plants and has been labelled with a taxon constraint of only being in the phylum Chordata, nuclear migration events have been observed in plants in processes like fertilization, shaping of elongated nuclei in root hairs, and plant-microbe interactions [36–38]. Indeed, the term “nuclear migration” (GO:0007097) has been assigned to genes in *Zea mays, Arabidopsis thaliana, Manihot esculenta*, and *Oryza sativa* in the GOA database. Upon further inspection, the GO term “interkinetic nuclear migration” (GO:0022027) is shown to be manually assigned only to a *Klebsormidium nitens* gene product (UniProtKB:A0A1Y1ICV9) in Viridiplantae in the GOA database. However, on the UniProt database [39], the annotation status of this protein seems to be unreviewed, and computer predicted. Since the GO terms in the GO Big plant subset were collected based on experimental and manual curation, this suggests that some gene annotations may need to be reviewed in terms of the type of evidence code assigned.

#### 3.3.2 Example 2: incorporation of GO terms of plant functions with non-inclusive ancestors

To investigate the reason behind the presence of the GO term “circulatory system development” (GO:0072359) in the GO Big plant subset, we also looked at the existence of any of its child terms in our subset. In fact, we were able to identify the GO term “regulation of vasculature development” (GO:1901342) as its most specific child term along the path of the DAG in the subset. In the GOA database, this GO term has been assigned to the *ABERRANT PANICLE ORGANIZATION* (*APO1* ) gene in *Oryza sativa* ssp. *indica* and *japonica* (UniProtKB:B8B183 and UniProtKB:Q655Y0). It has been reported that this gene plays a role in vascular bundle systems development [40]. Inspecting the parent terms of “regulation of vasculature development” (GO:1901342) shows that it has a “regulates” relationship with “vasculature development” (GO:0001944). Interestingly, although “vasculature development” is a biological process that does realistically exist in plants, it has been labelled with a taxon constraint of only being in the subphylum Vertebrata. In turn, “vasculature development” (GO:0001944) is labelled as “part of” the ancestor GO term “circulatory system development” (GO:0072359). Biologically speaking, the term “vasculature development” indicates a more species-inclusive process than the term “circulatory system development”. This indicates the presence of GO terms with ancestors representing less species-inclusive functions, making the higher-level terms less broad. This suggests that some high-level GO terms need to be reviewed either to become more taxon-neutral than their child terms or an alternate path in the DAG needs to be established with non-animal equivalent functions.

## 4 Discussion

### 4.1 The new GO Big DAG

We created a new type of GO subset, the GO Big subset, to improve the specificity of gene function prediction in plants. The main purpose of this type of subset is to contain biologically meaningful GO terms for one or more species of interest for accurate and precise prediction tools. Although slim GO subsets have been created and are often used, the purpose of GO Slim subsets is to give a generalized view of the functions represented in a group of genes for applications like high-throughput expression studies or annotation coverage evaluation [17, 41, 42]. The purpose of GO Big subsets – to get the most specific GO term possible without compromising its biological validity within a taxon - requires that the maximum possible specificity for the taxon be annotated. The biggest inspiration for the need of these GO Big subsets was the plant community. A common struggle in plant genomics is the annotation of some of the genes with errant functions. This problem is mainly due to the automatic propagation of these GO terms among genomes by computational pipelines [43].

As seen in the example detailed in our results, not only did both GO Big subsets produce more detailed GO descriptions for the maize gene *sdg130* as a histone methyltransferase compared to the GO Slim plant subset, but the originally assigned errant annotations were mostly substituted with more relevant and biologically valid functions within the maize species. However, although the GO Big subsets would present more specific plant function annotations than the GO Slim plant subset, there are some limitations that must be considered, as shown in the examples discussed in the previous section:

1. The GO Big subsets discussed in this study are an initial attempt to create a new type of species-specific and/or clade-specific subset. Although our subsets include taxon-neutral and plant-specific GO terms, creating a species-specific subset, like GO Big maize, can still lack common plant-specific functions and require expansion by including GO annotations from a broader clade to encapsulate diverse functions. The more species included, the better the coverage of plant functions, as seen between the GO Big maize and GO Big plant subsets.
2. The methods used to create these subsets are simple and user-friendly so that any researcher would be able to create one for their species of interest. However, this method needs to be coupled with manual expert and curatorial knowledge. For example, in the case of some plants, compared to their large number of genes, experiment-based annotations are scarce or nonexistant [44]. This phenomenon was observed for the species *Pinus lambertiana, Solanum pennellii, Vaccinium corymbosum L*., and *Coffea canephora*. Regions in the DAG consisting of GO terms describing uniquely plant processes, functions and components could be included in the GO Big plant subset, regardless of evidence codes. For example, the terms “extrinsic component of lumenal side of plastid thylakoid membrane” (GO:0035450) and “extrinsic component of stromal side of plastid inner membrane” (GO:0035454) clearly represent cellular components found in plants. Those terms were not annotated to any Viridiplantae gene - not even with an IEA evidence code - and were therefore not part of our subset. Additionally, the term “halotropism” (GO:0170002), a process that describes the tropic movement of roots in response to sodium [45], is also missing from our GO Big plant subset due to the lack of its assignment to any plant genes. Other examples of processes found in plants include “seed dehydration” (GO:1990068), “plant gross anatomical part developmental process” (GO:0160109), and “detection of parasitic plant” (GO:0002243). Perhaps, in this case, the availability of both an experimentally derived GO Big plant subset and another separate expanded GO Big plant subset that includes plant and taxon-neutral functions regardless of the evidence type may allow for uniquely useful applications in plant biology research.
3. There is an issue of research imbalance when it comes to genes of interest in plant and animal species [11]. Many researchers focus on studying genes of known functions when there still is a significant percentage of genes of unknown function in eukaryotic genomes [46]. Overcoming this bias may lead to the discovery of unique functions and the creation of new GO term annotations better suited to describe these novel processes, which could be electronically propagated to genes in other genomes, allowing the generation of new hypotheses to test, and ultimately leading to more experimentally verified annotations.
4. Another consideration is that the GO DAG itself (and therefore the GO Big plant subset) lacks good portrayal of diverse plant functions and may need to be expanded in some areas to include those unique plant biology aspects [13]. Most plant functions in the GO DAG were likely created for the model organism *Arabidopsis thaliana*. This single source for plant functions could lead to occurrences where the assignment of unusual functions is due to the absence of related plant functions in the GO graph. On the other hand, it is also worth noting that an unconventional function assigned to a plant gene may not be a false prediction, but simply one that has not yet been observed in plants or discovered experimentally.
5. As seen in this study, a few GO terms that appear errant were actually assigned to some plant genes manually, which resulted in their incorporation in our GO Big plant subset. Upon further inspection, these GO terms were mostly reflecting relationships, mainly regulatory interactions, between plants and other organisms. This underscores the importance of considering the context of the errant assigned annotation before excluding it from species-specific or clade-specific subsets. In addition, this suggests the possibility of missing GO terms from our subsets that describe such interactions due to the absence of assignments with experimental evidence codes, which again calls for the significance of curatorial expertise accompanying this method.
6. A way that we used to expand our GO Big plant subset was by considering taxon constraints, a correction system that includes the two relationships “only in taxon” and “never in taxon” to help curators flag inconsistencies in annotations [25] and is implemented in tools like the Phylogenetic Annotation Inference Tool (PAINT) and Argot2.5 web server [19, 25]. Indeed, this method was helpful for capturing GO terms that describe functions that are “only in” Viridiplantae, especially those that were not manually assigned to plant genes in the GOA database. On the other hand, not only were there some errant GO terms that were inherited in our GO Big plant subset that are ancestor terms of those describing plant functions as previously described in our results, but some of these ancestor terms were labelled with the “never in” Viridiplantae taxon constraint. This presented a problem as the removal of such terms would lead to the exclusion of entire areas in the DAG. For example, the exclusion of “epidermis development” (GO:0008544), which is labelled with the “never in” Viridiplantae taxon constraint, from our subset would lead to the subsequent exclusion of the child term “epidermal cell fate specification” (GO:0009957). This child GO term has been manually assigned to several genes (*GL2, BHLH2, GL3, TOP6A, TTG1*, and *WRKY44* ) found in *Arabidopsis thaliana* that play a role in determining the cell fate of root epidermal cells [47–51]. Using taxon constraints alone is not sufficient, and we suggest that these ancestor terms be revised to either become taxon-neutral or provide plant function-equivalent terms.

Future work entails updating the GO Big plant subset with newer file versions possibly by 1) collecting GO terms through curatorial knowledge regardless of evidence code, 2) including all taxon-neutral GO terms, and 3) creating subset variations in which GO terms from certain types of relationships are either included or excluded (as an example, choosing not to include GO annotations with “regulates” relations to avoid the integration of functions that describe interactions affecting other organisms).

### 4.2 How to GO Big

In this paper, we discussed creating GO Big with the ultimate goal to improve predicted gene function assignments in plants. By following this methodology, researchers can create GO Big subsets tailored to their specific taxon of interest. A practical application for these subsets is using them in tools like GOTermMapper [9, 31] to map errant GO terms, helping to reveal potential gene functions and inspire new hypotheses to test. An important feature for GO Big subsets is that, like GO, they are DAGs. This paves the way for a second application for GO Big: researchers and curators can identify areas of improvement within the graphs for missing or more inclusive function descriptions. Finally, a potential application of GO Big subsets is their integration into gene function prediction tools as an alternative to the full GO DAG, allowing researchers to obtain precise annotations relevant to the species’ biology without concerns about errant GO terms.

### 4.3 Why should you GO Big?

While the direct purpose of GO Big subsets is to obtain biologically meaningful annotations for the taxon of interest during gene function prediction, there is a larger implication. By restricting annotation assignments to functions valid to the biology of the organism, bias against the assignment of errant functions can be eliminated, allowing for a possible shift in research focus from highly-studied genes to poorly-characterized or unknown genes that have been electronically assigned with functions of interest. In other words, GO Big subsets may drive the advancement of science by indirectly addressing the problem of the significant number of unknown genes in different organisms. It is worth mentioning that considering gene function annotations derived from GO Big subsets along with the unconventional function predictions assigned to the same gene by the full GO DAG may spark new ideas about the possible function of the gene and inspire researchers to generate novel hypotheses to test, so even these annotations can be useful.

## 5 Conclusion

The abundance of sequenced plant genomes, the availability of the largest knowledgebase for gene functions, and the advancement in gene function prediction tools set up the right time and circumstances for the formation of the GO Big subset. In addition, the specific need for the GO Big plant subset arose from the comparatively lesser detail in knowledge of plant functions. By creating a new type of GO subset specifically curated for plants, we hope that this subset would be used by both function prediction tools and GO mapping tools to increase taxonomic unit specificity while restricting to biologically meaningful functions in plants. We also hope that the approach would be adopted and applied to other species resulting in more GO Big subsets. We expect the involvement of more biologists in generating such subsets would be critical in contributing to the GO Consortium’s mission in developing a comprehensive and inclusive model of biological systems.

## Supplementary information

Additional file 1.xlsx: Mappings of the 28 GO term annotations for the *Zea mays* B73v5 *sdg130* gene assigned by GOMAP using GOTermMapper. goslim plant: Excel sheet showing the resulting GO terms when the 28 GO annotations were mapped to the GO Slim plant subset. gobig maize: Excel sheet showing the resulting GO terms when the 28 GO annotations were mapped to the GO Big maize subset. gobig plant: Excel sheet showing the resulting GO terms when the 28 GO annotations were mapped to the GO Big plant subset. Rows in orange in the gobig maize and gobig plants sheets indicate that the terms are also present in the goslim plant sheet. Rows in blue in the gobig plant sheet indicate that the terms are also present in the gobig maize sheet, but not in the goslim plant sheet.

## Acknowledgments

We thank Colleen Yanarella, Leonore Reiser, Valerie Wood, Pascale Gaudet, and Chris Mungall for their helpful discussions. We also thank the Gene Ontology Consortium for taking the time and effort to create the Gene Ontology knowledgebase.

## Declarations

### Ethics approval and consent to participate

Not applicable

### Consent for publication

Not applicable

### Availability of data and materials

All data and source code generated are freely available at [26] under the terms of the MIT license.

### Code availability

All data and source code generated are freely available at [26] under the terms of the MIT license.

### Authors’ contributions

LF and CJLD designed the approaches to create the GO Big subsets. LF generated the GO Big subsets, organized the datasets, and performed data analysis. LF and CJLD wrote the manuscript. All authors read, suggested improvements, and approved the final copy of the manuscript.

## References

[1] Bard JBL, Rhee SY. Ontologies in biology: design, applications and future challenges. Nature Reviews Genetics. 2004 Mar;5(3):213–222. 10.1038/nrg1295.

[2] Ashburner M, Ball CA, Blake JA, Botstein D, Butler H, Cherry JM, et al. Gene Ontology: tool for the unification of biology. Nature Genetics. 2000 May;25(1):25–29. Number: 1 Publisher: Nature Publishing Group. 10.1038/75556.

[3] The Gene Ontology Consortium, Aleksander SA, Balhoff J, Carbon S, Cherry JM, Drabkin HJ, et al. The Gene Ontology knowledgebase in 2023. Genetics. 2023 May;224(1):iyad031. 10.1093/genetics/iyad031.

[4] Duan ZH, Hughes B, Reichel L, Perez DM, Shi T. The relationship between protein sequences and their gene ontology functions. BMC Bioinformatics. 2006 Dec;7(4):S11. 10.1186/1471-2105-7-S4-S11.

[5] du Plessis L, Škunca N, Dessimoz C. The what, where, how and why of gene ontology—a primer for bioinformaticians. Briefings in Bioinformatics. 2011 Nov;12(6):723–735. 10.1093/bib/bbr002.

[6] Wong ED, Miyasato SR, Aleksander S, Karra K, Nash RS, Skrzypek MS, et al. Saccharomyces genome database update: server architecture, pan-genome nomenclature, and external resources. Genetics. 2023 May;224(1):iyac191. 10.1093/genetics/iyac191.

[7] Jenkins VK, Larkin A, Thurmond J. Using FlyBase: A Database of Drosophila Genes and Genetics. In: Dahmann C, editor. Drosophila: Methods and Protocols. Methods in Molecular Biology. New York, NY: Springer US; 2022. p. 1–34. Available from: 10.1007/978-1-0716-2541-51.

[8] Ringwald M, Eppig JT, Kadin JA, Richardson JE, the Gene Expression Database Group. GXD: a Gene Expression Database for the laboratory mouse: current status and recent enhancements. Nucleic Acids Research. 2000 Jan;28(1):115–119. 10.1093/nar/28.1.115.

[9] Gene Ontology Consortium. The Gene Ontology (GO) database and informatics resource. Nucleic Acids Research. 2004 Jan;32(uppl 1):D258–D261. 10.1093/nar/gkh036.

[10] Gene Ontology Resource. Gene Ontology Resource Available from: http://geneontology.org/stats.html.

[11] Zhao Y, Wang J, Chen J, Zhang X, Guo M, Yu G. A Literature Review of Gene Function Prediction by Modeling Gene Ontology. Frontiers in Genetics. 2020 Apr;11. 10.3389/fgene.2020.00400.

[12] Wimalanathan K, Lawrence-Dill CJ. Gene Ontology Meta Annotator for Plants (GOMAP). Plant Methods. 2021 May;17(1). 10.1186/s13007-021-00754-1.

[13] Fattel L, Psaroudakis D, Yanarella CF, Chiteri KO, Dostalik HA, Joshi P, et al. Standardized genome-wide function prediction enables comparative functional genomics: a new application area for Gene Ontologies in plants. GigaScience. 2022 Jan;11:giac023. 10.1093/gigascience/giac023.

[14] Yanarella CF, Fattel L, Lawrence-Dill CJ. GWAS From Spoken Phenotypic Descriptions: A Proof of Concept From Maize Field Studies. 2023 Dec;10.1101/2023.12.11.570820.

[15] The Gene Ontology Consortium. The Gene Ontology Resource: 20 years and still GOing strong. Nucleic Acids Research. 2019 Jan;47(D1):D330–D338. 10.1093/nar/gky1055.

[16] Lomax J. Get ready to GO! A biologist’s guide to the Gene Ontology. Briefings in Bioinformatics. 2005 Sep;6(3):298–304. 10.1093/bib/6.3.298.

[17] Mutowo P, Bento AP, Dedman N, Gaulton A, Hersey A, Lomax J, et al. A drug target slim: using gene ontology and gene ontology annotations to navigate protein-ligand target space in ChEMBL. Journal of Biomedical Semantics. 2016 Sep;7(1):59. 10.1186/s13326-016-0102-0.

[18] Bilyk KT, Cheng CHC. Model of gene expression in extreme cold - reference transcriptome for the high-Antarctic cryopelagic notothenioid fish Pagothenia borchgrevinki. BMC Genomics. 2013 Sep;14(1):634. 10.1186/1471-2164-14-634.

[19] Lavezzo E, Falda M, Fontana P, Bianco L, Toppo S. Enhancing protein function prediction with taxonomic constraints – The Argot2.5 web server. Methods. 2016 Jan;93:15–23. 10.1016/j.ymeth.2015.08.021.

[20] Huntley RP, Sawford T, Mutowo-Meullenet P, Shypitsyna A, Bonilla C, Martin MJ, et al. The GOA database: Gene Ontology annotation updates for 2015. Nucleic Acids Research. 2014 Nov;43(D1):D1057–D1063. 10.1093/nar/gku1113.

[21] Binns D, Dimmer E, Huntley R, Barrell D, O’Donovan C, Apweiler R. QuickGO: a web-based tool for Gene Ontology searching. Bioinformatics. 2009 Sep;25(22):3045–3046. 10.1093/bioinformatics/btp536.

[22] Musen MA. The protégé project: a look back and a look forward. AI Matters. 2015 Jun;1(4):4–12. 10.1145/2757001.2757003.

[23] Jackson RC, Balhoff JP, Douglass E, Harris NL, Mungall CJ, Overton JA. ROBOT: A Tool for Automating Ontology Workflows. BMC Bioinformatics. 2019 Jul;20(1). 10.1186/s12859-019-3002-3.

[24] Klopfenstein DV, Zhang L, Pedersen BS, Ramírez F, Warwick Vesztrocy A, Naldi A, et al. GOATOOLS: A Python library for Gene Ontology analyses. Scientific Reports. 2018 Jul;8(1). 10.1038/s41598-018-28948-z.

[25] Deegan JI, Dimmer EC, Mungall CJ. Formalization of taxon-based constraints to detect inconsistencies in annotation and ontology development. BMC Bioinformatics. 2010 Oct;11(1). 10.1186/1471-2105-11-530.

[26] Fattel L, Lawrence-Dill CJ. Dill-PICL/gobig repository on GitHub; 2024. Available from: https://github.com/Dill-PICL/gobig.git.

[27] Batra R, Gautam T, Pal S, Chaturvedi D, Rakhi Jan I, et al. Identification and characterization of SET domain family genes in bread wheat (Triticum aestivum L.). Scientific Reports. 2020 Sep;10(1). 10.1038/s41598-020-71526-5.

[28] Zheng L, Ma S, Shen D, Fu H, Wang Y, Liu Y, et al. Genome-wide identification of Gramineae histone modification genes and their potential roles in regulating wheat and maize growth and stress responses. BMC Plant Biology. 2021 Nov;21(1). 10.1186/s12870-021-03332-8.

[29] Osadchuk K, Cheng CL, Irish EE. The integration of leaf-derived signals sets the timing of vegetative phase change in maize, a process coordinated by epigenetic remodeling. Plant Science. 2021 Nov;312:111035. 10.1016/j.plantsci.2021.111035.

[30] Lawrence-Dill CJ. Carolyn Lawrence Dill GOMAP Maize MaizeGDB B73 NAM 5.0 October 2022 v2.r1. CyVerse Data Commons; 2023. Available from: 10.25739/cfvb-jn16.

[31] Generic Gene Ontology Term Mapper; 2023. Available from: http://go.princeton. edu/cgi-bin/GOTermMapper.

[32] Kalervo Väänänen H. Mesenchymal stem cells. Annals of Medicine. 2005 Jan;37(7):469–479. 10.1080/07853890500371957.

[33] Xue W, Yu J, Chen W. Plants and Their Bioactive Constituents in Mesenchymal Stem Cell-Based Periodontal Regeneration: A Novel Prospective. BioMed Research International. 2018 Aug;2018:1–15. 10.1155/2018/7571363.

[34] Saud B, Malla R, Shrestha K. A Review on the Effect of Plant Extract on Mesenchymal Stem Cell Proliferation and Differentiation. Stem Cells International. 2019 Jul;2019:1–13. 10.1155/2019/7513404.

[35] Azizi A, Herrmann A, Wan Y, Buse SJ, Keller PJ, Goldstein RE, et al. Nuclear crowding and nonlinear diffusion during interkinetic nuclear migration in the zebrafish retina. eLife. 2020 Oct;9. 10.7554/elife.58635.

[36] Bone CR, Starr DA. Nuclear migration events throughout development. Journal of Cell Science. 2016 May;129(10):1951–1961. 10.1242/jcs.179788.

[37] Zhou X, Meier I. Efficient plant male fertility depends on vegetative nuclear movement mediated by two families of plant outer nuclear membrane proteins. Proceedings of the National Academy of Sciences. 2014 Jul;111(32):11900–11905. 10.1073/pnas.1323104111.

[38] Zhou X, Groves NR, Meier I. Plant nuclear shape is independently determined by the SUN-WIP-WIT2-myosin XI-i complex and CRWN1. Nucleus. 2015 Mar;6(2):144–153. 10.1080/19491034.2014.1003512.

[39] Bateman A, Martin MJ, Orchard S, Magrane M, Ahmad S, Alpi E, et al. UniProt: the Universal Protein Knowledgebase in 2023. Nucleic Acids Research. 2022 Nov;51(D1):D523–D531. 10.1093/nar/gkac1052.

[40] Terao T, Nagata K, Morino K, Hirose T. A gene controlling the number of primary rachis branches also controls the vascular bundle formation and hence is responsible to increase the harvest index and grain yield in rice. Theoretical and Applied Genetics. 2009 Nov;120(5):875–893. 10.1007/s00122-009-1218-8.

[41] Jaeger PA, Ornelas L, McElfresh C, Wong LR, Hampton RY, Ideker T. Systematic Gene-to-Phenotype Arrays: A High-Throughput Technique for Molecular Phenotyping. Molecular Cell. 2018 Jan;69(2):321–333.e3. 10.1016/j.molcel.2017.12.016.

[42] Rutherford KM, Lera-Ramíez M, Wood V. PomBase: a Global Core Biodata Resource—growth, collaboration, and sustainability. GENETICS. 2024 Feb;10.1093/genetics/iyae007.

[43] Wimalanathan K, Friedberg I, Andorf CM, Lawrence-Dill CJ. Maize G. Annotation—Methods, Evaluation, and Review (maize-GAMER). Plant Direct. 2018 Apr;2(4). 10.1002/pld3.52.

[44] Makrodimitris S, van Ham RCHJ, Reinders MJT. Automatic Gene Function Prediction in the 2020’s. Genes. 2020 Oct;11(11):1264. 10.3390/genes11111264.

[45] Szepesi Halotropism: Phytohormonal Aspects and Potential Applications. Frontiers in Plant Science. 2020 Sep;11. 10.3389/fpls.2020.571025.

[46] Wood V, Lock A, Harris MA, Rutherford K, Bähler J, Oliver SG. Hidden in plain sight: what remains to be discovered in the eukaryotic proteome? Open Biology. 2019 Feb;9(2):180241. 10.1098/rsob.180241.

[47] Ohashi Y, Oka A, Ruberti I, Morelli G, Aoyama T. Entopically additive expression of GLABRA2 alters the frequency and spacing of trichome initiation. The Plant Journal. 2002 Feb;29(3):359–369. 10.1046/j.0960-7412.2023.01214.x.

[48] Zhang F, Gonzalez A, Zhao M, Payne CT, Lloyd A. A network of redundant bHLH proteins functions in all TTG1-dependent pathways ofArabidopsis. Development. 2003 Oct;130(20):4859–4869. 10.1242/dev.00681.

[49] Schneider K, Wells B, Dolan L, Roberts K. Structural and genetic analysis of epidermal cell differentiation in Arabidopsis primary roots. Development. 1997 May;124(9):1789–1798. 10.1242/dev.124.9.1789.

[50] Galway ME, Masucci JD, Lloyd AM, Walbot V, Davis RW, Schiefelbein JW. The TTG Gene Is Required to Specify Epidermal Cell Fate and Cell Patterning in the Arabidopsis Root. Developmental Biology. 1994 Dec;166(2):740–754. 10.1006/dbio.1994.1352.

[51] Ishida T, Hattori S, Sano R, Inoue K, Shirano Y, Hayashi H, et al. Arabidopsis TRANSPARENT TESTA GLABRA2Is Directly Regulated by R2R3 MYB Transcription Factors and Is Involved in Regulation ofGLABRA2Transcription in Epidermal Differentiation. The Plant Cell. 2007 Aug;19(8):2531–2543. 10.1105/tpc.107.052274.

